# 3D kinematics of leaf-cutter ant mandibles: not all dicondylic joints are simple hinges

**DOI:** 10.1101/2023.08.28.555128

**Authors:** Victor Kang, Frederik Püffel, David Labonte

## Abstract

Insects use their mandibles for a variety of tasks, including cutting and material transport, defence, building nests, caring for brood, and competing for mates. Despite this functional diversity, mandible motion is thought to be constrained to rotation about a single fixed axis in the majority of extant species. Here, we conduct a direct quantitative test of this ‘hinge joint hypothesis’ in a species that uses its mandibles for a wide range of tasks: *Atta vollenweideri* leaf-cutter ants. Mandible movements from live restrained ants were reconstructed in 3D using a multi-camera rig. Rigid body kinematic analyses revealed strong evidence that mandible movement occupies a kinematic space which requires more than one rotational degree of freedom: at large opening angles, mandible motion is dominated by yaw. But at small opening angles, mandibles yaw and pitch. The combination of yaw and pitch allows mandibles to ‘criss-cross’: either mandible can be on top when mandibles are closed. We observed criss-crossing in freely cutting ants, suggesting that it is functionally important. Combined with recent reports on diversity of joint articulations in other insects, our results show that insect mandible kinematics are more diverse than traditionally assumed, and thus worthy of further detailed investigation.

## 1. Introduction

Jaws are an important evolutionary innovation for most vertebrates. Although the use of jaws for feeding is highly conserved, the variation in vertebrate jaw morphology and kinematics is remarkable [e. g. 1]. As illustrative examples, jaws in some teleost fish are integrated with other skeletal elements to render their skull into a complex multi-bar linkage [2]; mammals characteristically combine jaw translation and multi-axis rotation into complex chewing movements [e. g. 3–9]; and stingrays, salamanders, and some carps have convergently evolved jaw kinematics that resemble those observed in mammals [10–13]. The rich diversity and significance of jaw kinematics in vertebrates is reflected in the array of advanced quantitative techniques used to study them, including three-dimensional (3D) motion capture, X-ray reconstruction of moving morphology (XROMM), neural-network assisted automatic tracking, and rigid body mechanics [e. g. 9, 14–17].

In contrast, our quantitative and functional understanding of mandible movements in hexapods is nascent [e. g. 18, 19]. This discrepancy is partly owed to the experimental difficulties associated with capturing and digitising the motion of small and often fast-moving mandibles, and partly due to the need for high-resolution 3D imaging techniques such as lab-based computed micro-tomography, which have only recently become more accessible and widely used. Consequently, current perspectives on hexapod mandibular function have focussed on comparative morphology, qualitative in vivo observations, and detailed analyses of the evolutionary history of mandible morphology [20]. In the earliest hexapod lineages – the coneheads (Protura), springtails (Collembola) and two-pronged bristletails (Diplura) – mandibles have a single articulation point [21], and are actuated with multiple muscle groups [22–25]. However, in Dicondylia, a taxonomic group encompassing most extant insect species, the morphology of the mandible-head complex is markedly different: load-bearing structures of the head-capsules are strengthened [22, 26], mandibles articulate via two distinct condyles, and are typically actuated with one antagonistic pair of muscles [24, 25, 27–31]. For example, most of the winged biting-chewing insects that have been studied to date, including dragonflies [32], cockroaches [33], and beetles [34, 35], have dicondylic joints with two prominent mandibular condyles. Indeed, the transition from monocondylic to dicondylic mandibles is considered a pivotal moment in insect evolution [e. g. 23, 36, 37], paralleling the rise of jawed fish in early vertebrate evolution: it is thought to be associated with an increase in bite forces, and may have given early insects access to a wider range of food sources [22, 25, 37].

Another consequence of the emergence of dicondylic mandible joints is a putative simplification of possible mandible motion. A single condyle that articulates like a ball-and-socket joint allows three rotational degrees of freedom (DoFs). Two condyles, in turn, can form two ball-and-socket joints, which are thought to constrain mandible motion to rotation about a virtual axis which connects their centres [e. g. 24, 25, 31, 33]. Mandible joints of winged insects with biting-chewing mouthparts (Odonata and Neoptera) are thus classified as obligate dicondylic [25] – they are simple one DoF hinge joints which only permit rotation about a single fixed axis. In light of the staggering diversity and complexity of jaw movements in vertebrates, the alleged conformity to a simple hinge joint in the mandibles of the most speciose animal taxon may raise curious eyebrows. During the last 400 million years, insect mouthparts have undergone multiple independent and stunningly complex evolutionary modifications: to give but two out of many possible examples, in many hemipteran and dipteran species, biting mouthparts have been adapted into stylets for piercing, while in other Dicondylia, dicondylic joints have undergone a secondary reduction to return to monocondyly [20, 30, 31, 38]. Is it really plausible that all dicondylic mandibles of winged biting insects are adequately described by the same hinge joint paradigm? Although restriction to a single DoF simplifies neuromechanical control and may increase net bite force by channelling muscle force into two antagonistic muscle pairs [33, 35, 39–41], it is not free from disadvantages. In mammals, for example, multiple jaw DoFs enable an array of masticatory kinematics, with significant consequences for the ability to nutritionally process food [e. g. 42, 43]. Although many dicondylic mandibles possess regions that are ascribed a “grinding function” and likened to mammalian molars [e. g. 21, 24, 44–48], it would appear that two mandibles rotating about hinge joints are somewhat ill-suited for effective grinding, which requires relative motion between two surfaces that are pressed together.

Perhaps surprisingly, there is no quantitative evidence in support of the widespread idea that the kinematics of dicondylic mandibles in winged biting insects can be described by a single rotational DoF. One may argue that it is hard to imagine how a joint with two conspicuous condyles could depart from rotation about the axis they define. But few (if any) well-developed condyles form perfect ball-and-socket joints, and not all putatively dicondylic mandibles have two well-developed condyles. Indeed, recent work has revealed some cracks in the surface of the canonical assumption that all mandibles of winged biting insects rotate about a single fixed axis. Gronenberg et al., for example, inferred from high-speed video observations that mandible movements in a trap-jaw ant are suggestive of a mechanical “cam” within the joint, which is crucial for the successful preparation of a mandible strike [49]. This cam is formed by protrusions on the head-capsule, which guides a supposed sliding motion of the mandibular stem. Zhang et al. studied mandible kinematics in a ponerine ant, and suggested that bi-axial rotation of mandibles can confer multifunctionality: at small opening angles, ants can grasp delicate objects with concavities located on the ventral section of the mandible, but due to slight mandible roll, they can also cut and pierce prey with sharp teeth on the dorsal side when opening angles are large [50–52]. In several detailed studies on formicoid head morphology, it was noted that ant mandible joints may depart from a strict obligate dicondylic morphology, and instead have reduced or otherwise altered articulations [53– 55, see also [56]]. Using anatomical inference, computational visualisations and experiments with 3D printed models to indirectly infer mandible kinematics, it was suggested that here, too, the leading assumption of a simple hinge joint breaks down [53, 55]. Evidence for more complex articulation morphology is not restricted to ants: mandibles of parasitoid chalcid wasps appear to have reverted to “secondary monocondyly” to re-access DoFs that had been made unavailable by dicondylic joints [31].

Common to these studies is their use of morphological inference and qualitative observations as supporting evidence. Quantitative analyses of joint DoFs using rigid body mechanics, which are widely applied in vertebrates, are rarely used in insects [e. g. 57]. In this study, we address this gap, and use both qualitative and quantitative methods to characterise the mandible motion in *Atta vollenweideri* Forel, 1893 leaf-cutter ants. Leaf-cutter ants harvest plant fragments to cultivate a specialised fungus that serves as the primary food source for the colony [58, 59]. They forage on a wide range of plant materials, including leaves, flowers, and fruit of varying physical properties [60, 61]. Thus, leaf-cutter ants use their mandibles to bite and cut through materials that can be soft, hard, tough, thin, or thick, in addition to carrying out a variety of other tasks typical for ants, such as brood care, material transport, and nest building. We used marker-based motion capture to reconstruct mandibular motion of *A. vollenweideri* ants in 3D and *in vivo*, and then analysed the data using rigid body mechanics to investigate whether the observed kinematics are consistent with a simple hinge joint. These quantitative analyses are supplemented with descriptions of joint morphology, and observations of mandible movements during unrestrained cutting and biting to provide a holistic account of mandible joint function.

## 2. Methods

### (a) Study animals

A colony of *A. Vollenweideri* leaf-cutter ants, founded and collected in Uruguay in 2014, was used for all experiments. The colony was kept inside a climate-controlled chamber (FitoClima 12.000 PH, Aralab, Rio de Mouro, Portugal), on a 12/12 hour day/night cycle at 25°C and 60% relative humidity. It had access to leaves of bramble, laurel, and kibbled maize *ad libitum*, regularly provided in two foraging boxes connected to the colony via plastic tubing.

### (b) Behavioural observations, manual mandible manipulation and scanning electron microscopy

Ant workers cutting either bramble leaves or polydimethylsiloxane polymer films (PDMS; approximate thickness: 300 μm) were video-recorded using a Raspberry Pi High Quality camera (Raspberry Pi Foundation, Cambridge, UK), equipped with a 25 mm C-mount lens (LM25JC, Kowa Optimed Deutschland GmbH, Düsseldorf, Germany), and a Raspberry Pi model 3B. For observations of mandible movements at higher magnifications, individual worker ants were secured in a 3D printed mount and recorded under a stereomicroscope (Z6, Leica Microsystems GmbH, Wetzlar, Germany). Thick mid-veins of bramble leaves were offered to the worker ants to observe mandible movements involved in cutting.

Three mature workers, between 19–20 mg in body mass, were selected from the foraging box connected to the colony, and sacrificed by freezing. Immediately after isolating the ant head using a scalpel, the mandibles were carefully manipulated under a stereomicroscope (S Apo, Leica Microsystems GmbH, Wetzlar, Germany), to qualitatively assess their range of motion. The head-capsule was held with tweezers, and a second pair of tweezers was used to gently push the right mandible downwards as both mandibles were fully closed. One manipulation trial was filmed with a smartphone through the eye-piece of the stereomicroscope (Google Pixel 6a, 1920*×*1080 pixels, 30 frames per second (fps)).

Following manual manipulation, one of the three head samples was halved using a scalpel. The left hemisphere was left intact, but the mandible from the right hemisphere was removed by holding it near the distal end with tweezers. All samples were air-dried in a desiccator for four days, and then mounted on aluminium stubs, using carbon tape and silver conductive paint, to prepare them for scanning electron microscopy (SEM). The specimens were sputter-coated with approximately 30nm of gold (Emitech K575X, Quorum Technologies, Laughton, UK), and imaged in secondary electron mode using a tungsten filament SEM (JSM-6010LA, JEOL Ltd., Tokyo, Japan).

#### Multi-camera rig for 3D motion tracking

To characterise *in vivo* mandible motion in 3D, a multi-camera recording rig was constructed with 30 *×* 30 mm aluminium struts and 3D printed joineries (Fig. 1 a). Four CMOS USB cameras (2*×* model DMK 33UX265, 2*×* DMK 33UX252; The Imaging Source Europe GmbH, Bremen, Germany), each equipped with a Telecentric lens (Computar TEC-M55, CBC America, NC, USA), were mounted on this rig via custom-designed camera mounts with five degrees of freedom (DoF), which helped to adjust camera orientation and lens focus. The object of interest – either a live specimen or a calibration target – was first mounted on a 3D printed platform, which was then slotted into a 3D printed stage with three DoF, used to position the sample within the field of view of the cameras. The scene was illuminated with LED lamps (LED desk lamp, IKEA, Delft, The Netherlands), and the cameras were synchronised using a custom-built external trigger comprising an Arduino Nano microprocessor, programmed to send pulse signals to the trigger port of each camera in parallel. Synchronised images were captured at a resolution of 2048*×*1536 pixels, and at 60 fps, using the software provided by the camera manufacturer (IC Capture v2.5.1547.4007, The Imaging Source).

**Figure 1.**
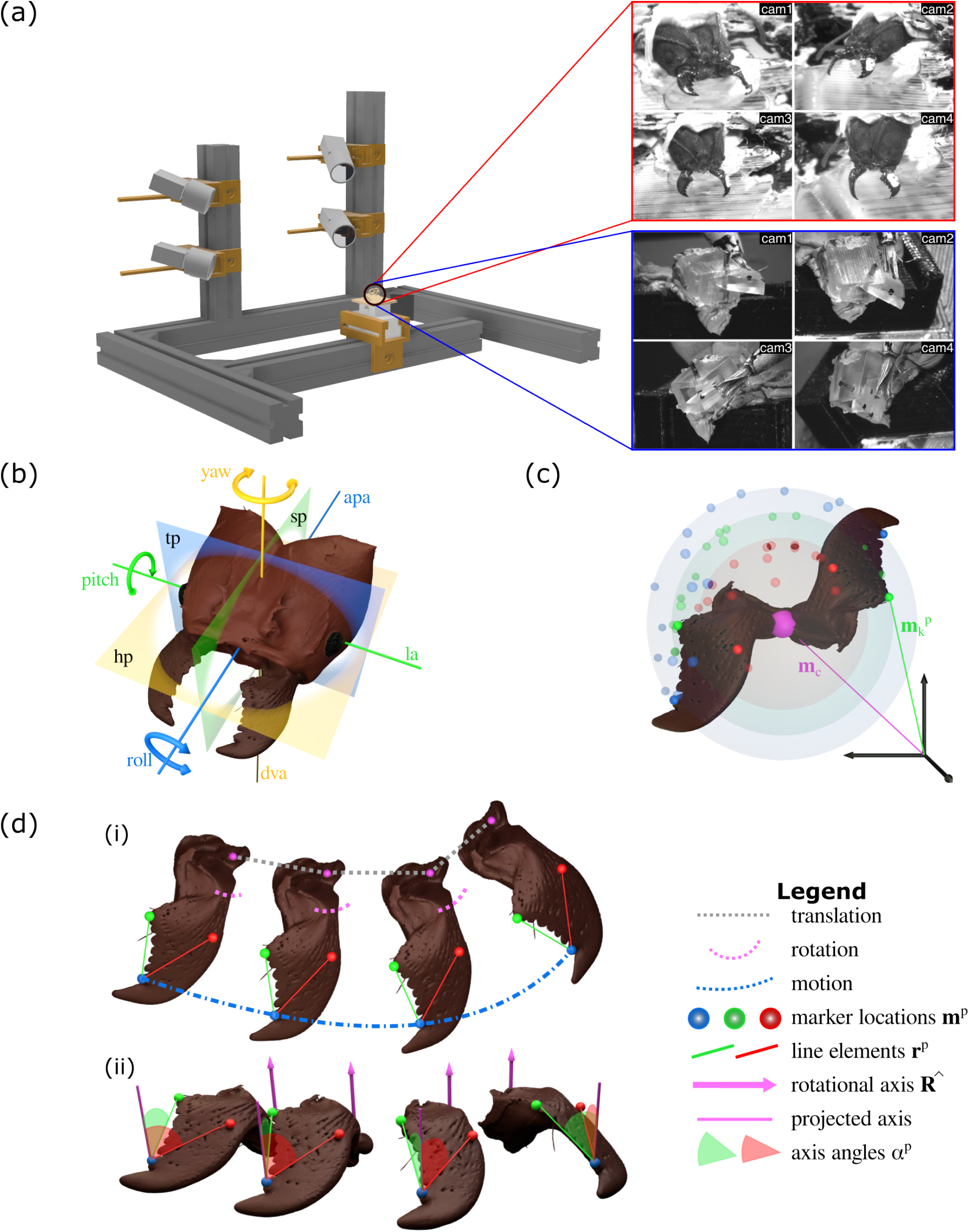
3D motion capture and principles of quantitative kinematic analysis of mandible motion in *Atta vollenweideri* leaf-cutter ants. (a) A four-camera recording rig built from aluminium struts and 3D printed joineries was used to record mandible motion in 3D. Individual ants were restrained on a moveable holder (black circle), and direct linear transformation coefficients were estimated for 3D reconstruction. Top red inset: example views of a live restrained worker ant. Three markers on each mandible were used for tracking, and are visible as white spots: one distal (blue sphere in (c-d)), one intermediate (red sphere in (c-d)), and one proximal (green sphere in (c-d)). Bottom blue inset: example views of a physical hinge joint model - built from 3D printed parts and an insect pin - used as a reference for quantitative analyses. (b) Schematic of an *A. vollenweideri* head-capsule defining three principal planes — sagittal (sp), transverse (tp), and horizontal (hp); three principal axes — dorso-ventral (dva), anterior-posterior (apa), and lateral (la); and three principal rotations — pitch, yaw, and roll. The hypothesis that *Atta* mandible joints are hinges was tested in two steps. (c) First, if mandibles rotate about a fixed point (pink), then all markers trace surfaces of concentric spheres. The location of the best joint centre, **m**_**c**_ was thus estimated by minimising the variation of the distances between all three markers locations 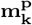 and the joint centre across all frames (see eq. 2.2). For a hinge joint, the distance variation must be negligible. (d-ii) Second, if mandibles rotate about an axis with fixed orientation, then any line element connecting two points on the mandible spans a constant angle to this axis throughout the rotation, independent of mandible or joint centre translation. The orientation of the best continuous axis of rotation (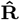, pink) was thus estimated by minimising the variation of the angles *α*^*p*^ spanned by it and two mandible-fixed line elements **r**^**p**^ across all frames (red and green lines and angles, see eq. 2.3 and main text for details). For a hinge joint, the difference between the predicted and observed angle should thus be close to zero, and uncorrelated, continuous, independent and normally distributed with respect to the angle of rotation about 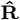.

A 1*×*1 LEGO®plate was used as calibration target, a common choice for 2D and 3D calibration due to LEGO®’s strict manufacturing standards [e. g. 62, 63]. Real-world distances between nine landmarks on the calibration target were measured with a digital micrometer (Model 293-795-30, Mitutoyo Corporation, Sakado, Japan; maximum permissible error *±*2 μm). Synchronised images of the target were then taken on all four cameras. The images were loaded into a MATLAB (R2021b, v9.10.0; The MathWorks Inc., Natwick, MA, USA) application to estimate the direct linear transformation (DLT) coefficients for camera calibration [64], which requires identification of at least six landmarks; all the visible landmarks from the calibration object (n=9 total landmarks) were identified from each of the four camera views. The camera calibration was validated ahead of every experimental trial involving a re-positioning or a new specimen: the current calibration was loaded into DLTdv8a [64, 65], along with a short image sequence from each of the four camera views. The location of the apical tooth tip of the left mandible was selected in one random playback window, and the corresponding 2D projection line was inspected in the three remaining views. If the 2D projection lines were in satisfying agreement, the same location was selected in a second playback window, which then returned the 3D projection and the DLT residuals. A system was considered well-calibrated if the DLT residuals were below 1.0 pixels; otherwise, it was redone.

#### Video recording, tracking, and 3D reconstruction

For each trial, one *A. vollenweideri* worker was selected from the foraging box. We selected six ants within a narrow body mass range to minimise potential size effects (20.4–23.6 mg). Each ant was positioned on a custom 3D printed mount which featured a clamp, designed to physically restrain the head-capsule during the recordings. The mounted specimen was chilled on an ice block for 1-2 minutes to reduce activity during subsequent marker placement. Markers were added to enable consistent and accurate tracking, using a small insect pin dipped in correction fluid (Tipp-Ex Rapid, Société Bic S.A., Paris, France). Two constraints informed marker placement: to increase the accuracy of downstream kinematic analyses, the marker positions needed to be far away from each other and yield non-collinear line elements, at least one of which is approximately perpendicular to a perceived dominant axis of rotation (see below); and markers needed to remain visible on at least three cameras throughout the movement. Markers were added to the second and last tooth (defined as most proximal), as well as at a coordinate near the mid-point of the mandible, opposite to the masticatory margin of the left mandible (Fig. 1 a & c). The mounted and marked ant was kept on the ice block until the set-up was ready for acquisition; this helped to reduce active mandible movements during the wait period, which often led to loss of the markers via mandible scraping. Preliminary trials indicated that the brief cooling period had no discernible impact on subsequent behaviour. Some ants readily displayed biting movements, so the cameras were triggered, and around 2000 – 3000 frames were captured (corresponding to 30-50 s recording time); otherwise, individuals were encouraged to move their mandibles by a gentle puff of air. Ants used for the trials were released back to the foraging box after filming was completed.

All recordings contained multiple mandible movement sequences, and were reviewed post-trial in ImageJ v1.53j [66]. Frame sequences that contained mandible movements were exported and loaded into DLTdv8a [64], along with the calibration file, for marker tracking. Six landmarks were manually located on each frame: one on the approximated volumetric centre of each eye, one halfway between the ventral mandibular articulation (*vma*) and the atala (*ala*), to approximate the mandible joint location [morphological terms as per 67], and three points on the marked mandible as described above. The output of the tracking was a 3D position vector for each landmark per frame.

### (c) Kinematic analysis

The extracted position vectors were used for 3D kinematic analyses, with the aim to test the hypothesis that mandibles of leaf-cutter ants are connected to the head via hinge joints, and thus rotate about a single axis fixed both in space and in orientation. Characterisation of 3D joint kinematics is a common aim in skeletal biomechanics, and particularly in human medicine and sports science [e. g. 68–72]. In classic rigid body kinematics, a general analysis would proceed by first characterising the orientation and then the location of an instantaneous helical axis (*IHA*) – the axis about which the body rotates and along which it translates at a given instant. The orientation of the *IHA* can be calculated exactly, through use of the rigid body constraint: all relative motion between two points on the same rigid body is due to rotation, and all points on the body share the same instantaneous angular velocity vector, ***ω***; by definition, ***ω*** has the same orientation as the *IHA* [73, Throughout this manuscript, we denote vectors as bold; unit vectors are indicated with a hat, as in 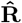 for a unit vector in the direction of the vector **R**.]. Three non-collinear points, *A, B*, and *C*, on the rigid body then suffice to determine ***ω*** via a direct vector solution of a set of two simultaneous equations, which reads [68]:

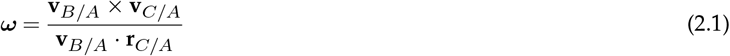

where **v**_*B/A*_, **v**_*C/A*_ and **r**_*C/A*_ are the instantaneous relative velocity and relative position vectors for points *A, B* and *C*, respectively. The location of the *IHA* can then be found relative to any reference point *X* on the body via an expression which again involves the angular velocity and the instantaneous linear velocity of point *X* [73]. This two-step process is both theoretically exact and general, and is consequently widely used [e. g. 72, 74–79].

In principle, rotational joint DoFs can then be inferred by inspecting the temporal variation of the orientation and location of the *IHA* throughout a motion sequence. For a hinge joint, for example, the *IHA* degenerates to a single rotational axis with constant orientation and location. However, the instantaneous relative and absolute velocity vectors, required to estimate the orientation and location of the *IHA*, are generally unknown, and thus need to be approximated via the average relative velocities over a finite time, equal to the instantaneous velocities only in the limit of an infinitesimal time step [see e. g. 75, for a more detailed discussion]. This approximation introduces a direct competition between two needs: the time step between two measurements needs to be as small as possible, so that the average velocity vectors approach the instantaneous velocity vectors; but it needs to be large enough to avoid vanishing signal-to-noise ratios. The strength of eq. 2.1 is thus also its weakness: because it is an exact vector solution, any noise is necessarily interpreted as a change in the orientation and location of the *IHA*.

The sensitivity of the instantaneous helical axis to noise is a well-known problem, and can be so severe that determination of its orientation and location can be prohibitively inaccurate, even for the much simpler case of planar motion, where it reduces to an instantaneous axis of rotation [e. g. 69, 75, 80]. Indeed, even relatively small noise can reduce the estimate obtained through eq. 2.1 to no better than a random guess (see supplementary information, SI). To overcome this difficulty, numerous alternative methods have been proposed. These methods may be broadly split into one of two categories: the orientation of the *IHA* is still obtained through determination of the angular velocity vector, but filtering or other optimisation routines are deployed to reduce noise [e. g. 69, 75, 80]; or finite difference approximations are avoided altogether, and a representative axis is estimated through minimisation of cost functions which allow absorption of experimental uncertainty into an error term [e. g. 71, 77, 81–83]. Although these methods can reduce the sensitivity to noise, they are typically restricted to rotation about a fixed point or axis [77, 81]. Other approaches exist, including some which allow joint axes translation [e. g. 74, 82, 83], but a comprehensive account of these methods and a rational comparison of their individual merits and shortcomings is beyond the scope of this study.

Because leaf-cutter ant mandibles are small (length of about 1.8 mm), we anticipated that even best-case signal-to-noise ratios may be such that direct calculation of the angular velocity vector would yield noise-ridden estimates of the *IHA*, preventing robust conclusions on rotational joint DoFs through inspection of its temporal variation. In order to conduct a direct test of the hinge joint hypothesis in *Atta* mandibles, we instead developed a method which leverages the advantages of cost functions, but can estimate the orientation of the rotational axis independent of joint translation. The general idea is to estimate the axis with fixed orientation and location which describes as much of the observed kinematics as possible – we will call this axis the best continuous axis of rotation, *bCAR*. The hypothesis of a dicondylic hinge joint will then be assessed quantitatively by statistical comparisons of the explained versus observed mandible kinematics.

In order to estimate the location of the *bCAR*, we define a cost function that minimises the variation of the distance between the location of *P* markers, 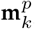, and the best joint centre of rotation, **m**_*c*_, across *N* frames [see 81]:

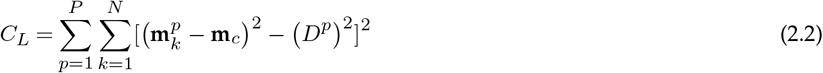

Here, *D*^*p*^ are the average distances between each **m**^*p*^ and **m**_*c*_ across all frames. If the mandible rotates about a fixed point, then all markers populate the surfaces of concentric spheres with radii *D*^*p*^, and minimisation of Eq. 2.2 yields the coordinate of their shared centre [Fig. 1 c. See ref. 81, for a more detailed discussion]. However, a unique solution exists only if rotation occurs about more than one axis; if rotation is constrained to a single axis of rotation, *any* point on it minimises eq. 2.2, because all points now populate the circumference of concentric circles. In order to obtain a point that is biologically meaningful, we thus restricted the possible coordinate of **m**_*c*_ such that it falls within in a square bounding box with a side length of 20 % of the distance between both eye centres (about 600 μm), around the putative joint centre (see above).

In order to determine the orientation of the rotational axis, 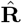, we introduce the elemental geometric notion that the angle between two Euclidean vectors, **r** and 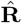, can be defined via their dot product as cos 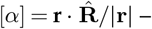 the basis of the widely used Rodrigues’s rotation formula [84]. Let **r** be an arbitrary vector which connects two points on a rigid body, and let this body rotate by a movement angle *θ* about an axis with fixed orientation 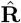 . The angle *α*, and thus 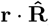, remain constant throughout the rotation. More generally, any line element **r**^**p**^ which connects two points on a rigid body forms a constant angle *α*^*p*^ to 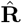 throughout any rotation about 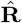 (Fig. 1 d). Crucially, this angle is invariant to translation of either the body or the rotational axis. 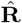 can thus be estimated via a cost function which minimises the variation of *α*^*p*^ for *P* − 1 independent vectors spanned by *P* markers across *N* frames:

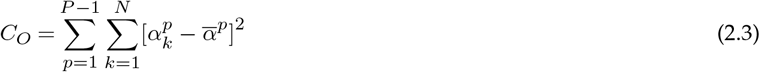

where 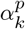 is the angle between 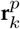 and 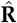 at the *k*th time instant, and 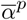 is the arithmetic mean angle for each vector:

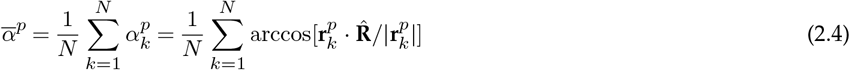

In a sense, the cost function defined by eq. 2.3 is a reformulation of eq. 2.1, with the key difference that it can consider data from an arbitrary number of line elements to fit a single axis to an arbitrary number of observations, as opposed to determination of one single solution per point pair via a finite difference approximation. Thus, for *P* = 3 markers and *N* = 2 frames close in time, minimisation of eq. 2.3 yields an estimate for the orientation of the *IHA*; but in practise, it is then subject to similar trade-offs as laid out above (see SI for a more detailed performance evaluation).

To apply eq. 2.3 to our experimental data, we defined two line elements **r**^*p*^ for each mandible, as the vectors connecting the most distal marker to the other two markers. All 3D kinematics data per individual were then subjected to the cost function defined by eq. 2.3 to determine the orientation of the *bCAR* per individual.

### (d) Statistical assessment of the hinge-joint hypothesis

In order to test how well mandible kinematics can be described as rotation about a fixed point, we assessed the variation of the distance between **r**^*p*^ and **m**_*c*_ across all recorded frames per individual. The magnitude of this variation was then assessed by comparing it to (i) the accuracy of the distance measurements on the calibration target (*±*2 μm, see above); (ii) the size of one image pixel (7 μm); (iii) the lengths of the mandible (about 1.8 mm) and of the effective mandible in-lever [about 600 μm, see 41]; and (iv) the variation obtained for a physical model hinge joint (see below and Fig. 1 a).

To test whether mandible kinematics can be accurately described as rotation about an axis with fixed orientation, we extracted the residuals resulting from the minimisation of eq. 2.3 – the difference between expected and observed angles 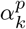 – and then inspected their distribution across the movement angle range *θ*_*max*_. The movement angle range *θ*_*max*_ was defined as the largest angle that was spanned between all vectors which connect the *bCAR*, located through eq. 2.2, with the most distal marker in the plane of rotation. If mandible motion is restricted to rotation about a single axis with fixed location and orientation, these residuals should be small, continuous, independent and normally distributed with respect to the movement angle *θ*; *θ* can be defined with respect to any reference vector that connects the *bCAR* to the most distal marker in the plane of rotation, and we defined it such that it is zero when mandibles were fully opened. To formally assess these necessary statistical conditions, we conducted an ordinary least squares regression between residuals and movement angle *θ*, and then subjected the residuals to the Global Validation of Linear Models Assumptions test developed by Peña and Slate[85], implemented in the R-package *gvlma*, using the global test statistic with a significance level of 0.05. The null hypothesis of rotation about an axis with fixed orientation was retained only if there was no significant relationship between the residuals and movement angle, and if the linear model assumptions were met. Two further parameters were extracted as proxies for the quality of fit: the standard deviation of the residuals resulting from minimisation of the cost function (averaged across both line elements); and the ratio between the observed and predicted distance moved by the mandible, from here on referred to as distance metric. The observed distance is the total distance covered by the individual markers; the predicted distance is the sum of distances between the marker locations projected onto the plane of rotation.

In order to compare results across ants, we converted individual-specific movement angles into global opening angles independent of the orientation of the rotational axis; we defined the opening angle as the angle spanned by the vector which connects the joint centre with the most distal tracking point and the normal of the sagittal plane (Fig.1 b). All kinematic calculations were conducted in python v3.9.13, and all statistical analyses were conducted in R v4.1.2. All data reported in the text correspond to mean *±* standard deviation unless otherwise indicated.

#### Validation

In order to validate the experimental and analytical methods, we tested them on experimental data obtained from a well-defined physical model with a hinge joint (inset in Fig. 1 a). The model, consisting of a ‘head-capsule’ and a ‘mandible’, was designed in Autodesk Fusion 360 (Autodesk, Inc., San Rafael, CA, USA) to have similar dimensions to a 20 mg worker ant head-capsule and mandible, and was printed using a stereolithography resin printer (Form3, Formlabs, Somerville, MA, USA). The mandible was held in place within the head-capsule by an insect pin that served as rotational axis; rotation about this axis was induced with a gentle push from a fine-tip paintbrush (n=3 instances from one model). The resulting movement was recorded using the set-up described above, and the 3D coordinates of the following landmarks were extracted for each frame: three points on the mandible (one distal, one intermediate, one proximal); two points on the longitudinal axis of the insect pin to serve as the ground truth rotational axis; and two points on the head-capsule to define a horizontal vector orthogonal to the rotational axis. The data were then subjected to the same analysis as described above to estimate the error associated with the 3D reconstruction and tracking, and the accuracy of the rigid body kinematic analysis.

#### Computed microtomography and 3D visualisation

3D printed models and computer animations were used to intuitively explore and visualise mandible kinematics [see also 55]. To this end, we used a computed micro-tomography scan of an *A. vollenweideri* worker with a body mass of 23.7 mg, obtained for a previous study [86], and similar in body mass to the workers investigated herein. The surface meshes of mandibles and head-capsule, segmented previously in ITK-SNAP [86], were exported out of ITK-SNAP as OBJ files and decimated using the ‘quadratic edge collapse simplification’ function in MeshLab v.2020.03 [87]. Meshes were further cleaned in MeshLab, and then imported into Blender v2.91.0, an open-source 3D animation software. Mandibular motion was visualised based on either (1) the real-world tracked points, or (2) the *bCAR* and the opening angle derived from the same points, using data from representative worker of similar body mass (21.1 mg).

## 3. Results

### (a) Mandible joint morphology

The mandible interfaces with the head-capsule via the mandible stem. The stem bears the ventral mandibular articulation (*vma*), the atala (*ala*), and the dorsal mandibular articulation (*dma*), which together define the joint (see Fig. 2 a). The *vma* and *ala* resemble ball-and-socket joints; the *dma*, in contrast, is elongated and akin to a hinge (Fig. 2 a & b). The cuticular surface of the *ala* is partially covered in a dense field of short spine-like protuberances, which terminate in either a single peak or multiple peaks (Figure 2 a-iii). A region of the *ala* surface that is microscopically smooth and completely devoid of protuberances is exposed when the mandible is fully closed (Fig. 2 a-ii-iii). Long slender hairs are present on the lower (ventral) half of the *al*, but not on its dorsal half (Fig. 2 a-iii). Both the *vma* and the *dma* cuticular surfaces lack the spine-like protuberances. The slender hairs are present, but with lower density (Fig. 2 a-iii).

**Figure 2.**
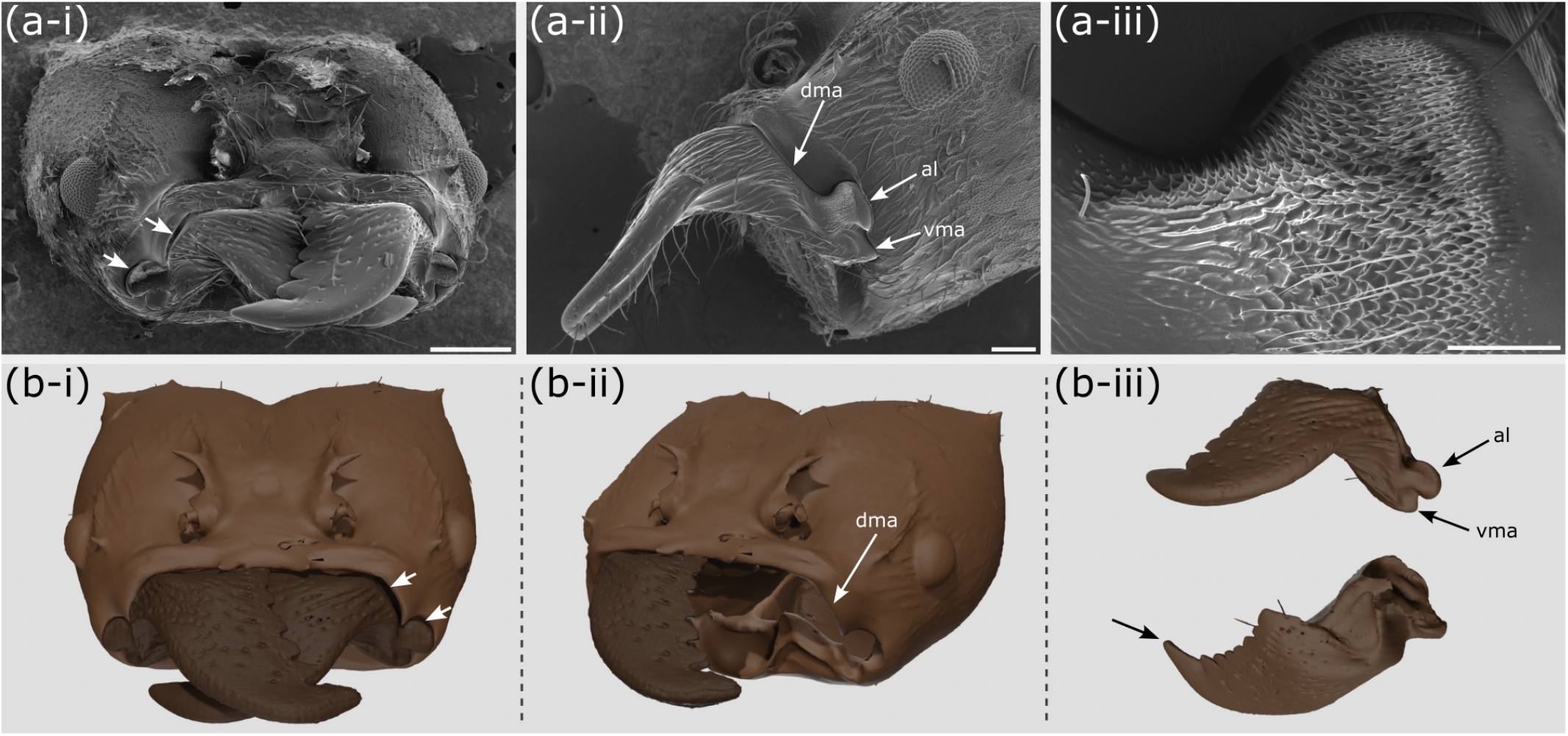
Morphology and kinematics of *Atta vollenweideri* leaf-cutter ant mandibles. (a-i) Scanning electron microscopy image of a worker ant head-capsule. Closed mandibles overlap, and one mandible must consequently be inferior (right in this case). For the inferior mandible, there is a clear gap between the head-capsule and the dorsal mandibular articulation (white arrows), which is larger than the gap in the same region of the superior mandible. Scale bar: 0.5 mm. (a-ii) Lateral view of the head-capsule, the left mandible, and the mandible joint. The joint morphology is defined by the dorsal mandibular articulation (*dma*), the atala (*ala*), and the ventral mandibular articulation (*vma*). Scale bar: 0.2 mm. (a-iii) The surface of the *ala* is covered in spine-like protrusions and hairs. The surfaces of the *dma* and *vma* are smooth. Scale bar: 0.05 mm. (b-i) 3D rendering of the head-capsule and left mandible of a different *A. vollenweideri* specimen, generated from computed micro-tomography images (false colours). White arrows indicate the gap between the inferior left mandible and the *dma*, also evident in (a-i). (b-ii) An ant head-capsule was digitally dissected to remove the left mandible and muscles. The *dma* is elongated and distinct in shape from the *vma* and *ala*. (b-iii) Lateral and dorsal views of the left mandible (top and bottom, respectively). The arrow marks the apical tooth on the masticatory margin.

### (b) Qualitative observations of mandible kinematics during cutting and biting

When cutting bramble leaf lamina, *A. vollenweideri* worker ants draw their mandibles from approximately fully open to almost fully closed. This motion visually resembles transverse adduction (Fig. 3 a and Supplementary Video (SV) 1). Mandibles were sometimes closed sufficiently to overlap, in particular when cuts were initiated at the free leaf edge, or when workers attempted to cut thick leaf veins (Figure 3 b and SV 2). Notably, there was no apparent pattern of preference for either the left or right mandible ending up as superior. Indeed, we observed swaps in superiority *in vivo* during cutting of thick leaf veins, leaf lamina, and thin PDMS films (Fig. 3 b and SV 2 & 7). That both the left and right mandible can be superior suggests that the kinematic space accessible to the mandibles may not be limited to rotation about a fixed axis: mandible “criss-crossing” likely requires mandible pitch (rotation in the sagittal plane), in addition to yaw (rotation in the horizontal plane, Fig. 1 b). We also observed this pitch movement in experiments with restrained live ants (Fig. 3 c & d): when a worker ant was mounted such that her head was held steady and the left mandible was physically prevented from opening, the mandible moved downward upon interaction with the right mandible (Fig. 3 c). When the right mandible re-opened, the left mandible sprung back to its initial position (Fig.,3 c-iv). This joint elasticity was retained in a freshly dissected ant head: when a gentle torque was applied to a closed mandible using tweezers, it pitched approximately about the horizontal axis of the *vma*, before returning to the original position upon removal of the torque (Fig. 3 d-i to iv and SV 3) – a motion visually resembling the pitch observed in live ants (Fig. 2 d-iv; SV 4). Thus, the mandible joint articulation appears to permit pitch in regions of mandible overlap.

**Figure 3.**
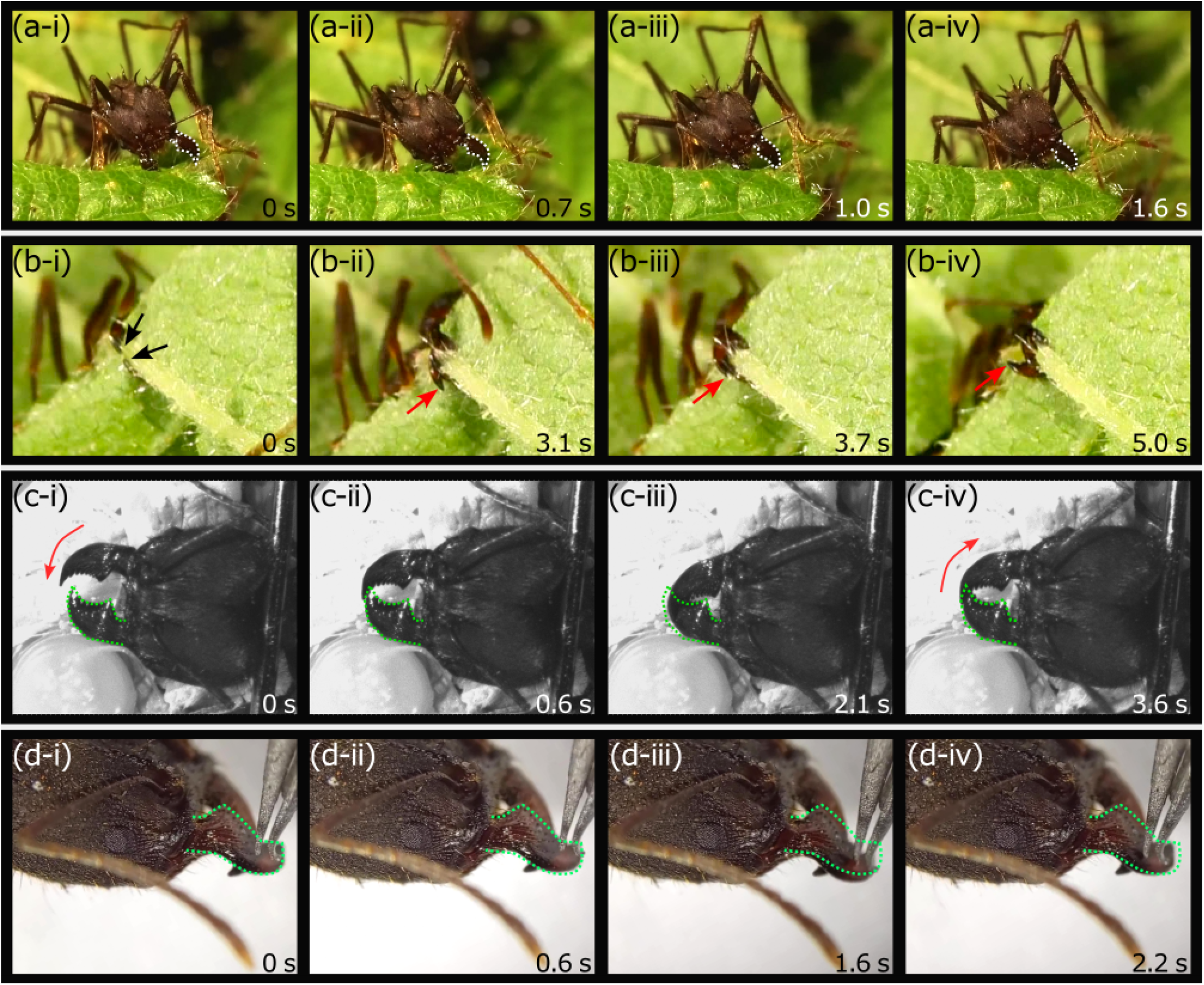
Qualitative observations of *Atta vollenweideri* mandible movement. (a-i to iv) A worker ant uses her left mandible (dotted outline) to slice through bramble leaf lamina, which dominantly involves mandible yaw (transverse adduction, SV 1). (b-i to iv) A worker ant is cutting through a thick bramble leaf vein, viewed from below. The black arrows label the apical tooth of each mandible; the red arrows highlight the inferior mandible. In b-ii and b-iii, the right mandible is inferior, and in b-iv, the left mandible is inferior, indicating that mandible yaw was combined with pitch during these closing movements (SV 2). (c-i to iv) Mandible motion in a live but restrained worker. The left mandible (green outline) was physically prevented from opening, while the right mandible was free to move at will. From (c-i) to (c-iii), the right mandible is superior. The position of the left mandible as seen in (c-i) is imprinted in dotted green onto (c-i) to (c-iv). In (c-iii), the left mandible temporarily pitches downwards, and then returns to its previous position once the right mandible is no longer on top in (c-iv), again illustrating that the mandible articulation permits mandible pitch when mandibles are fully closed. (d-i to iv) A similar motion of the mandible can be reproduced by manual manipulation of a dissected worker ant head. Gentle application of a downward force with tweezers caused the right mandible (green outline) to pitch downwards (d-i to iii). Upon removal of the force, the mandible rapidly returned to its initial position, as indicated again by the dashed green outline (d-iv; also refer to SV 3). Refer to Fig. 2 to infer approximate length-scales (all ants used were of similar size).

### (c) Quantitative analysis of mandible kinematics

The 3D kinematic space occupied by mandible movements mirrored the qualitative observation of mandible movements during unrestrained cutting and biting: mandible motion is dominated by transverse adduction, but mandibles appeared to both pitch and yaw in regions of mandible overlap (see illustrative examples in Fig. 4 a & b and SVs 5 & 6). In order to formally assess whether the overall motion patterns can be explained by rotation about a fixed axis, we subjected kinematic data from six *A. vollenweideri* specimens to quantitative kinematic analysis.

**Figure 4.**
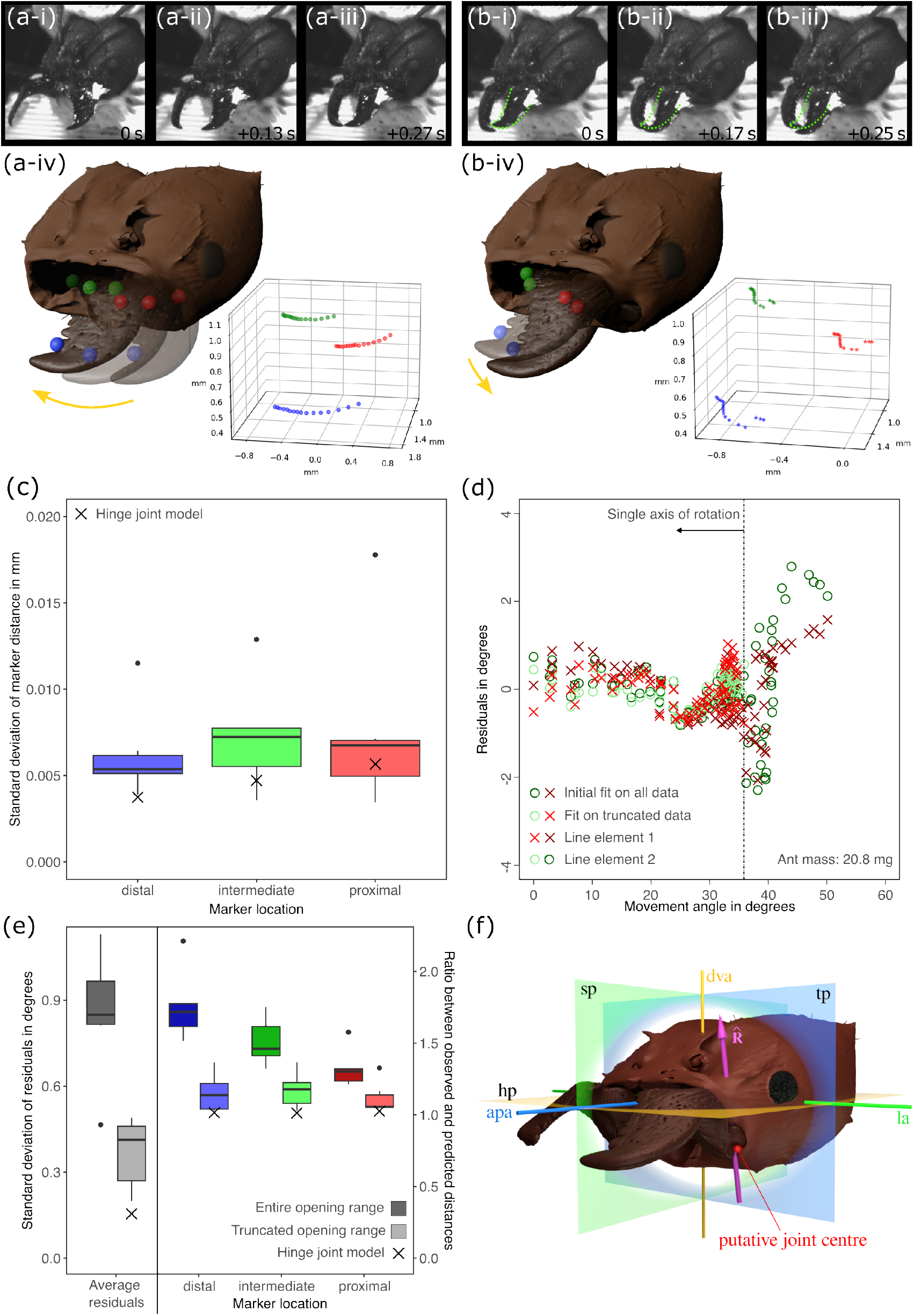
Quantitative analysis of *Atta vollenweideri* mandible motion. Mandibles displayed one of two characteristic movement patterns, mirroring the qualitative observations reported in Fig. 3: (a) transverse adduction and abduction, i. e. mandible yaw; and (b) a combination of mandible yaw and pitch, which occurred exclusively in regions of mandible overlap. (a-i to iii) Illustration of mandible yaw. (a-iv) 3D rendering and 3D kinematics of mandible yaw (ant mass: 21.1 mg). (b-i to iii) Illustration of mandible pitch and yaw. The initial position of the left mandible is highlighted in green. (b-iv) 3D rendering and 3D kinematics of mandible pitch and yaw (ant mass: 21.1 mg). Coloured spheres represent the position of three tracked markers. (c) Quantitative kinematic analysis of data from six individuals confirmed that mandible movement is consistent with the expectation of rotation about a fixed point: The distances between markers and the estimated joint centre of rotation vary little across the kinematic space, and the average distance variation is comparable to the base line variation observed for a physical hinge joint model with comparable dimensions (crosses). (d) The variation of the angles formed by two line elements on the mandibles and the best continuous axis of rotation statistically violates the expectation for uniaxial rotation. Qualitative inspection of the data suggested that this violation arises from a widening of the residuals in the region of mandible overlap (large movement angles), reflecting heteroscedasticity (see Supplementary Information for plots from all individuals). (e) To account for this observation, we estimated the largest movement angle range statistically consistent with uniaxial rotation. This analysis split the kinematic space into two characteristic regions (see main text for details): for small movement angles (large opening angles), mandible kinematics were consistent with the hinge joint hypothesis (truncated range). The standard deviation of residuals and the distance metrics were significantly reduced, and close to data obtained from a physical hinge joint model (crosses). In contrast, the statistical assumptions were violated for data from large movement angles (small opening angles), indicating that rotation can be multi-axial in the region of mandible overlap (not shown). (f) The estimated dominant axis of rotation 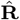 (pink) for large opening angles was closely aligned to the dorso-ventral head axis (dva); the location and orientation of this axis was highly consistent across individuals, and close to the initial guess informed by morphology (red sphere).

To test the hypothesis that ant mandibles rotate about a fixed point, we estimated the location of a joint centre as the coordinate which minimises the variation in the distances between the joint centre estimate and mandible markers across the kinematic space (see eq. 2.2). A suitable way to assess the hypothesis is to inspect the variation of these distances: 1.62 *±* 0.006 mm, 1.07 *±* 0.007 mm and 1.17 *±* 0.008 mm for the distal, intermediate and proximal marker, respectively (between 90 and 185 distances for each of n=6 ants, Fig. 4 c). The standard deviation of these distance indicates negligible deviation from rotation about a fixed point: it corresponds to about three times the accuracy of the calibration target dimensions, about one image pixel, less than 1% of the mandible length, less than 5% of the effective in-lever, and is in good agreement with the standard deviation obtained from a physical hinge joint model, which serves as a proxy for experimental noise (3.526 *±* 0.004 mm, 2.361 *±* 0.005 mm, 2.311 *±* 0.006 mm, respectively). Furthermore, the derived joint centre was in close proximity to the initial guess informed by morphology (putative joint centre): it was shifted toward the sagittal plane by 0.067 *±* 0.037 mm, or 5-10 pixels (n=6, Fig. 4 f). To test the hypothesis that ant mandibles rotate about a single axis with fixed orientation, we estimated the orientation of the axis that minimises the variation of the angle spanned with two mandible-fixed line elements across the kinematic space — the orientation of a best continuous axis of rotation (*bCAR*, see eq. 2.3). For all but one ant, the residuals of at least one line element violated the assumptions of the regression analysis, indicating that a single axis cannot accurately explain the observed kinematics (see SI for detailed plots for all individuals; n=6). Qualitative inspection of the residuals suggested that this result emerged from a systematic widening of the residuals in the region of mandible overlap, hinting at heteroscedasticity (Fig. 4 d). In contrast, for widely open mandibles, the residuals were small and consistently close to the expected zero line (Fig. 4 d & e). In order to account for this systematic pattern in our analysis, we next determined the largest movement range for which the residuals were uncorrelated, continuous, independent, and normally distributed. To this end, we iteratively truncated the kinematic space to smaller and smaller movement ranges, starting from the region of mandible overlap where residuals appeared inflated. We then re-calculated the *bCAR*, residual angles, and the movement angle range after each truncation step, and re-tested the residual angles until all assumptions were met for both line elements (Fig. 4 d and SI; Type 1 error rate was controlled via the Holm–Bonferroni method). This procedure suggested that mandible kinematics were consistent with rotation about a single fixed axis as long as opening angles were larger than 68 *±* 9°; for reference, mandibles start to overlap for opening angles smaller than about 80°, and displayed a global minimum and maximum opening angle of 49 and 114° across all ants, respectively. The cut-off angle is thus consistent with the qualitative observations in freely cutting and biting ants, which suggested that mandibles can both pitch and yaw in the region of mandible overlap (Fig. 3 b and SV 2). The average cut-off angle was subsequently used to split the kinematics space of each individual into two characteristic regions. For each truncated region, the *bCAR*, the standard deviation of residuals, and the distance metric were extracted separately, and compared to the global values across the entire kinematic space.

For opening angles larger than *>* 68°, which encompass most of the mandible movement range, the inference that a single axis of rotation sufficiently explains the mandible kinematics was supported further by two independent pieces of evidence (Fig. 4 e). First, both the standard deviation of the angle residuals and the distance metrics for all three markers decreased significantly (0.85 *±* 0.22° vs 0.37 *±* 0.13°; paired t-test: t_5_ = 7.17, p *<* 0.001. Distal distance metric: 1.76 *±* 0.24 vs 1.16 *±* 0.13; paired t-test: t_5_ = 7.73, p *<* 0.001; intermediate distance metric: 1.51 *±* 0.16 vs 1.17 *±* 0.12; paired t-test: t_5_ = 3.9, p *<* 0.05; proximal distance metric: 1.32 *±* 0.13 vs 1.12 *±* 0.11; Wilcoxon rank sum test: W = 21, p *<* 0.05. n=6 for all tests). To put both values into perspective, the corresponding results for the physical hinge joint model were 0.16 ° and 1.02, 1.01 and 1.03 for the distal, intermediate and proximal distance metrics, respectively (Fig. 4 e). Second, the orientation of the *bCAR* was highly consistent across individuals, with a mean deviation about the global average of 3.0 *±* 1.0° (n=6, Fig. 4 f). In all individuals, the rotational axis was approximately aligned with the dorso-ventral head axis (deviation of 6 *±* 2°), and thus lies dominantly in the sagittal and transverse head planes (deviation of 4 *±* 2° for both. Fig. 4 f). Given that the global orientation estimate is influenced by tracking error, fitting error, inaccuracy in head landmark placement, and biological variation, the small deviation across individuals strongly suggests a robust estimate of a consistently oriented *bCAR* in this region of the kinematic space.

In contrast, for opening angles smaller than 68°, the residuals of all but one ant still significantly violated the assumptions of the regression analysis for at least one line element (see SI for plots for all individuals; n=6). Thus, rotation in this region is multi-axial, consistent with the initial speculation based on qualitative observations of mandible kinematics. In further support of this conclusion, both the standard deviation and the distance metrics in this region increased upon truncation (1.05 *±* 0.26° for the standard deviation, and 2.59 *±* 0.53; 1.88 *±* 0.36; and 1.52 *±* 0.22 for the distance metric of the distal, middle, and proximal marker, respectively); the markers moved a distance between 1.5 and 2.5 times larger than predicted for rotation about one fixed axis.

## Discussion

The majority of winged insects with biting-chewing mandibles are thought to exhibit obligate dicondyly: they have mandible joints that act as simple hinges, permitting only rotation about a single fixed axis. This supposed kinematic simplicity remains quantitatively unconfirmed, and stands in marked contrast to jaw movements in vertebrates, which often involve translation and multi-axial rotation. We used 3D motion capture and rigid body mechanics to conduct a direct test of the hinge-joint hypothesis in *A. vollenweideri* leaf-cutter ants. Mandible kinematics were consistent with rotation about a single fixed axis across most of the kinematic space, but deviated from simple hinge joint kinematics around the region of mandible overlap. Hence, contrary to the long-held view that the dicondylic joints of winged biting-chewing insects are hinges, mandibles kinematics in leaf-cutter ants involve more than one degree of freedom.

### (d) The orientation of the dominant axis of rotation controls bite performance, and mandible yaw, roll and pitch

The majority of the kinematics space occupied by mandible movements reflects mandible yaw, and can be accurately described as rotation about a single dominant axis of rotation. This conclusion is supported by three independent lines of evidence. First, the distance between mandible markers and a derived joint centre is highly consistent across the kinematic space (Fig. 4 c); second, fitting a single rotational axis to data from opening angles larger than 68° resulted in small and randomly distributed residuals of the cost function defined by eq. 2.3 (Fig. 4 e); and third, the standard deviation of the residuals and the distance metrics in this region of the kinematic space were small, and comparable to those obtained from a physical hinge joint model (Fig. 4 e).

The orientation of the dominant axis defines a unique plane of rotation, and thus determines the magnitude of the mechanical advantage of the force transmission system [25, 33, 35, 41], the possible mandibular gape with respect to the head- or body fixed anatomical planes [25, 33, 41], and the angle between apodeme and effective in-lever [41]. All these parameters play a crucial role in determining insect bite performance [41]. All previous work in insects implicitly or explicitly assumed that the mandible rotational axis has a fixed orientation and location, so that a unique instantaneous mechanical advantage can be defined [25, 33, 35, 40, 41, 86, 88, 89]. Our results in *A. vollenweideri* indeed suggest that the rotational axis has an approximately fixed position, but for two reasons, this should only be a minor source of comfort for those interested in exact mechanical models of insect bite performance. First, joint articulations in insects are distinct from idealised shapes such as spheres or cylinders, and there is no strong reason to postulate *a priori* that the location of the rotational axis is generally fixed in space. Vertebrate joints virtually never have an instantaneous rotational axis which is fixed in space, and there is little reason to assume this is consistently different in insects. Second, even seemingly negligible movements of the joint centre can cause significant errors in mechanical calculations. As an illustrative example, a 2 cm misplacement of the human hip joint centre results in a decrease of the transmittable muscle moment of close to 50 % [90]. The key problem is that insect skeletal segments are small, so that highly accurate 3D data is required to resolve relevant displacements. It is conceivable that variation of the mechanical advantage due to small movements of the joint centre explains some of the residual variation in *Atta* bite force that cannot be captured by mechanical models which assume a fixed joint centre location [41].

The functional implications of the rotational axis for bite mechanics are well appreciated, but the behavioural and functional significance of its orientation with respect to a head- or mandible-fixed coordinate system has received less attention [25, 50–52]. In early hexapods, 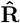, appears to be closely aligned with the distal incisivi and is nearly orthogonal to “molar areas” [25]. Mandibles dominantly rotate about their long axis, and this rolling motion may aid grinding, but limits the possible mandible gape [25]. In silverfish (Zygentoma) and winged insects (Pterygota), 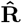 shifted; it is oriented more towards the cephalo-caudal head axis in orthognathous and hypognathous insects [25], or the dorso-ventral head axis in prognathous insects. Consistent with this general trend, the dominant rotational axis in prognathous *A. vollenweideri* deviates by less than 6° from dorso-ventral axis. Mandible rotation thus largely involves yaw, i. e. contraction of the mandible opener and closer muscles results in transverse mandible abduction and adduction, respectively. In contrast to the roll of early hexapod mandibles, mandible yaw enables large mandible gapes [25], which is essential for ants that use their mandibles to grasp objects comparable in size to their head width, such as brood, or, more specific to the leaf-cutters, to cut large fragments from leaf laminas (Figure 4 ai-iv and SV 3). However, alternative orientations may bring other functional benefits. In general, if the rotational axis has components in more than one of the three body planes, the mandible opening angle will have components in all three planes – mandibles will yaw, roll, and pitch with respect to head-fixed reference planes. Thus, bi-axial mandible movement with respect to head or body planes, as observed in a trap-jaw and a ponerine ant [49, 50], is not conclusive evidence against hinge joint kinematics *per se*. Instead, it may merely suggest an inclined rotational axis, advantageous because it may allow ants to handle objects of different sizes with different parts of their mandibles [50–52].

### (e) The origin and functional significance of mandible pitch

Qualitative observations of mandible movements during cutting and biting suggested that mandibles can pitch: both the left and right mandible can be superior when mandibles are closed, and swaps in position occurred across repeated closing cycles (Figure 3 & 4 and SV 4 & 7). Quantitative evidence for pitch was extracted from 3D kinematics data: for opening angles smaller than 68°, the residual angles unexplained by the *bCAR* varied systematically with the movement angle (see SI figure), and the mandible markers moved distances considerably larger than expected from rotation about a fixed axis (see Fig. 4 e). Thus, our results provide strong evidence against the null hypothesis of a simple hinge joint in *A. vollenweideri* ants. How is mandible pitch actuated, and what is its biological function?

Each mandible in *A. vollenweideri* is actuated by a single pair of antagonistic muscles. Each muscle is attached to the mandible via its own apodeme, and thus has one main line of action [86]. Activation of these muscles results in rotation or translation of the mandible, unless the net muscle force and moment are balanced by reaction forces and moments sustained by the joint articulation (or by the reactions associated with bites into a stiff object). Because the reaction sustainable by the joint depends on the joint articulation, and because the shape of this articulation may vary with opening angle [see e. g. 53, 55], it is conceivable that the sustainable reactions vary as the mandible rotates, so permitting multi-axial orientation in restricted sections of the kinematic space. Thus, even a simple antagonistic muscle pair can in principle drive complex rotation and translation. In *Atta*, the net muscle vector is almost perfectly aligned with the anterior-posterior axis [41, 86], and can thus induce both yaw and pitch moments. Perhaps pitch is blocked by the articulations at large mandible gape, but becomes possible in the region of mandible overlap, as indicated by the presence of a gap between the *dma* and the head capsule for the inferior mandible (Fig. 2). Alternatively, pitch may result from the interaction of the two mandibles: the reaction at the contact point between both mandibles could re-direct the muscle force, enabling a combination of yaw and pitch. Indeed, we did not observe deviations from rotation about the *bCAR* when mandibles were far away from each other, which is consistent with, if not sufficient to conclusively support, both hypotheses. Pitch of overlapping mandibles was also recently observed in *Odontomachus brunneus* trap-jaw ants: during rapid mandible closure, mandibles first followed a circular trajectory, but out-of-plane movements occurred as mandibles overlapped [91]. Re-direction of muscle forces via reaction forces between mandibles, or a variation of sustainable joint reactions due the changes in articulation shape, may be two valuable strategies to enable more complex “mastication-like” kinematics of dicondylic mandibles, without the need to resort to more complicated and energetically costly direct actuation via different muscle groups.

Based on our observations of both unrestrained and restrained cutting, we suggest three functional benefits of mandible pitch. First, leaf-cutter ants frequently use their mandibles like scissors, for example when they initiate cuts at the free edge of a leaf-lamina [92, 93]. As any hairdresser can attest, scissors need to be tight to function, and in man-made scissors, this is typically ensured by a screw which connects two blades, each with a slight inward curvature, at a shared pivot point. As the scissors are drawn close, the blades bend, and the stored elastic strain energy helps to keep the gap between the blades small, and the scissors tight [94]. In *A. vollenweideri*, a similar effect may help to keep the blades tightly connected during scissor-cutting. Indeed, in both living and dead specimens of *A. vollenweideri*, the gap between the two mandibles was narrow (Figs. 2 a-i & 4 b), and mandibles of dead ants sprung back to their original position after pitch was induced by application of an external torque, indicating joint elasticity. Second, during the cutting of tough materials such as primary or secondary leaf-veins [95], mandible criss-crossing may help to find a path of least resistance, and thus reduce the mechanical effort for cutting. Indeed, mandible criss-crossing and slight re-orientations of the mandibles were often observed when ants made repeated attempts to cut through thick leaf veins [Fig. 3 b and supplementary material in 92], reminiscent of how we may attempt to cut a thick wire with pliers or a thick tree branch with loppers. Third, mandible pitch may prevent damage to the sharp cutting edge during mandible impact. In one behavioural observation, a worker ant was cutting through a thin polymer sheet, which bend easily and unpredictably (SV 7). When the ant attempted to close its mandibles, the sheet flexed, and the left mandible went rapidly from superior to inferior and back.

#### Mandible kinematics and joint morphology in Dicondylia are more diverse than traditionally assumed

We have demonstrated that *A. vollenweideri* mandible joints have more than one degree of freedom. Mandible joints with multiple DoFs were likely the original condition in early hexapods, which possessed monocondylic mandibles actuated by multiple muscle groups [25]. Silverfish and firebrats (Zygentoma), jumping bristletails (Archaeognatha), and mayflies (Ephemeroptera) have retained a complex muscular actuation, but mandible motion appears to be more tightly constrained by a set of non-permanent articulations [22, 24, 25], which may allow for “gliding movements” in addition to rotation [20, 22]. These taxa thus share a more complex mandible muscle actuation with early monocondylic hexapods, but possess more restrictive mandible joint articulations reminiscent of obligate dicondyly – a trait combination that has consequently been referred to as “facultative dicondyly” [22]. In Odonata and Neoptera with biting mouthparts, muscle actuation reduced to a single pair of antagonistic muscles, and the consensus is that two conspicuous condyles restrict mandible kinematics to rotation about a single axis; mandibles are obligately dicondylic [25]. Remarkably, a reversion to a more complex muscle actuation and less restrictive joint articulations appears to have occurred in a group of parasitic wasps (Chalcidoidea), which was reported to have a strongly reduced posterior condyle and a modified musculature [31]. These modifications presumably enable active actuation of multiple DoFs [31]. In light of its resemblance to the ancestral trait of early hexapods, this mandible joint configuration has been referred to as “secondary monocondyly” [31]. In *A. vollenweideri*, joint articulations also depart from classic dicondyly: the ventral articulation (*vma*) resembles a ball-and-socket joint, but the dorsal articulation (*dma*) is an elongated, cylindrical, and thus conspicuously distinct from the *vma* in shape (Fig. 2 a). Without the coupling of dorsal and ventral ball-and-socket joints to constrain the mandible – as observed in obligate dicondylic taxa – additional DoFs become accessible. On the basis of these observations, it is evident that neither facultative dicondyly, obligate dicondyly, nor secondary monocondyly adequately describe the mandibles of *A. vollenweideri*. Instead, we propose that they represent “secondary facultative dicondyly,” characterised by the following conditions: two condyles exist, but have distinct morphology; (2) mandible kinematics involve more than one DoF, but the dominant movement is rotation about a single fixed axis, actuated by a single antagonistic pair of muscles; and (3) additional DoFs emerge through changes in sustainable joint reaction throughout the kinematic space, or via physical interactions between overlapping mandibles. Recently, an elongated *dma* has been identified as a characteristic trait of the Formicidae [54, 96], raising the possibility that secondary facultative dicondyly may be widespread in ants.

Jaw kinematics in vertebrates have received significant attention, but the equivalent problem in insects remains quantitatively unexplored. There is an increasing body of evidence which suggests that insect mandible kinematics are considerably more diverse than traditionally appreciated – representatives of at least two insect families that were thought to have simple hinge joints are capable of more complex mandible movements. Why did some neopteran taxa shift away from obligate dicondyly? Perhaps the more diverse mandible movement repertoire of wingless hexapods confers functional advantages that remain under-appreciated [25]. Quantitative comparative studies are necessary to unravel the evolutionary pathways and driving factors that lead to a reversion to more complex mandible movements in some insects, and to understand their functional consequences.

## Supporting information

Supplementary information raw data Atta wollenweideri

Supplementary information raw data physical model

## Acknowledgements

We thank Prof. Flavio Roces for his generous support, and for gifting the *Atta vollenweideri* colony used in this work, Prof. Tyson Hedrick for rapid replies and valuable advice on using DLTdv8a, and Prof. Alexander Blanke for stimulating discussions on insect mandible joints. This study is part of a project that has received funding from the European Research Council (ERC) under the European Union’s Horizon 2020 research and innovation programme (Grant agreement No. 851705) and a Human Frontier Science Programme Young Investigator Award (RGY0073/2020), both to DL.

