## supplementary_information for "3D kinematics of leaf-cutter ant mandibles: not all dicondylic joints are simple hinges"

Victor Kang*, Frederik Püffel* and David Labonte

Department of Bioengineering, Imperial College London

**Synthetic data and error analysis**

In order to assess the accuracy of our rigid body analysis, we computer-generated sets of kinematic data. Three markers were placed in a 3D coordinate system such that they formed three non-collinear relative position vectors; the maximum distance to the origin for each marker was set to $l_{max} = 1$. For each marker, circular trajectories were then generated by rotating the respective location vectors around a single predefined rotational axis, $\hat{\mathbf{R}} = (0, 0, 1)$, using Rodrigues' rotation formula (Rodrigues, 1840). The trajectories consisted of 30 equidistant time points (frames, $N$), and span an angular range of 60° (similar to the experimental values obtained in this study, see main text). Three-dimensional noise was then introduced to the locations of each marker in order to investigate the robustness of different methods to estimate $\hat{\mathbf{R}}$. Noise was incorporated by drawing a pseudo-random number from a normally-distributed population around zero, which was then added to each coordinate (see Gamage and Lasenby, 2002). The standard deviation of this normal distribution, $\sigma_{noise}$, was kept constant across markers and trajectories, but was varied linearly between 0 and $0.03\, l_{max}$ to generate independent datasets with varying levels of noise (31 noise levels with 50 repetitions each, see Fig. S 1). Translation of the rotational axis was not considered, as our cost function based on the minimisation of an angle difference is independent of translation (see main text).

For each noise level and repetition, we first determined the orientation of the instantaneous helical axis, *IHA*, via eq. 2.1 in the main manuscript, as done in traditional rigid body kinematics. We then subjected the data to the cost function defined by eq. 2.3 in the main manuscript to determine the best continuos axis of rotation, *bCAR* across all frames for each single simulation. Because the *bCAR* is estimated via minimisation in a numerical fitting routine, an initial orientation guess is required. In order to avoid positive bias, we selected an unfavourable initial guess, which spanned an angle of $\approx 80°$ to the true axis, $\hat{\mathbf{R}}$. To evaluate the accuracy of each method, we then calculated the angle, $\varepsilon_{angle}$, span by the estimates for $\hat{\mathbf{R}}$ and its true orientation.

For noiseless data, all three methods yield exact results, i. e. $\varepsilon_{angle} = 0$ as expected (see Fig. S 1a). However, even for small levels of noise, this exercise suitably demonstrated the difficulty associated with attempts to infer joint degrees of freedom from the temporal variation of the *IHA* (e. g. Ancillao et al., 2022; Stokdijk et al., 1999; Woltring et al., 1985): the *IHA* varies considerably between adjacent frame pairs (see purple vectors in Fig. S 1b and c). Frustratingly for the experimentalist, this problem gets worse if the spatio-temporal resolution is increased, due to diverging signal-to-noise ratios (see main text and Woltring et al., 1985). One way to overcome this problem is to average the pairwise *IHA* across all frames, to then use this estimate in a residual analysis as suggested in the main manuscript. However, even the global average of the *IHA* performs poorly compared to the *bCAR* in the presence of noise (see Fig. S 1d), and is no better than an average guess ($\varepsilon_{angle} \approx 45°$) for intermediate noise levels of $\sigma_{noise} = 0.015\, l_{max}$ – a region in which the *bCAR* is still accurate to better than 10°. Alternatively, the *IHA* may also be estimated for each possible frame pair, and then averaged. Although this procedure yields the most accurate estimate (see Fig. S 1d), it is plagued by statistical and conceptual concern: it is reminiscent of pseudo-replication, and it is anything but obvious how any deviation from rotation about this axis could be associated with a specific frame (or angular region) in residual analyses, as each frame is linked to numerous $(N - 1)$ instantaneous axes of rotation. Clearly, further analysis is required to assess the comparative benefits of these different methods, and to determine their accuracy for a broad range of tracking data.

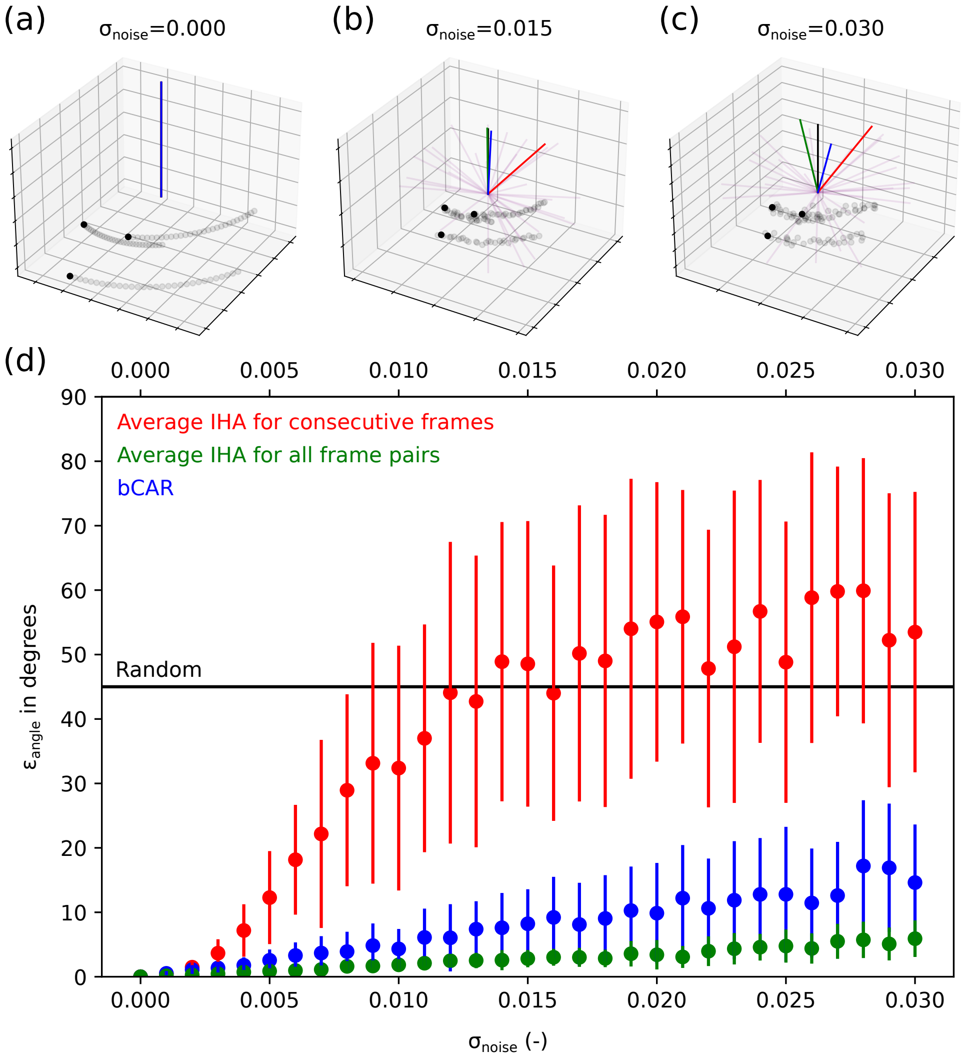


Figure S 1 | **(a-c)** We compared the accuracy of two ways to estimate the orientation of the rotational axis: via the instantaneous helical axis *IHA* (purple), as done in traditional rigid body kinematics; and via the cost function defined by eq. 2.3 in the main manuscript (blue). To assess the performance of each method, we computer-generated kinematic data. Each dataset contained a total of 90 position vectors from three markers, which were rotated about a pre-defined axis of rotation (black) across 60°. Varying levels of Gaussian noise were added to the marker locations, characterised by a standard deviation ($\sigma_{noise}$) between 0 and 3% of the maximum distance between tracked points and the axis of rotation, $l_{max}$. For noiseless data, either method is exact, as expected. However, even for small noise, the *IHA* varies considerably across frames, rendering inference of joint degrees of freedom all but impossible. **(d)** One way to overcome this problem is to calculate a representative average *IHA* across all frames. However, even this more robust estimate performs worse than the *bCAR* for all but the smallest noise levels, and is reduced to no better than a random guess as noise exceeds 0.015 $l_{max}$. An average *IHA* could also be estimated as the average *IHA* across all possible frame pairs (green). Although this procedure yields the most accurate estimate, it is associated with conceptual and statistical problems (see text).

**Rodrigues, O.** (1840). Des lois géométriques qui régissent les déplacements d'un système solide dans l'espace, et de la variation des coordonnées provenant de ces déplacements considérés in-

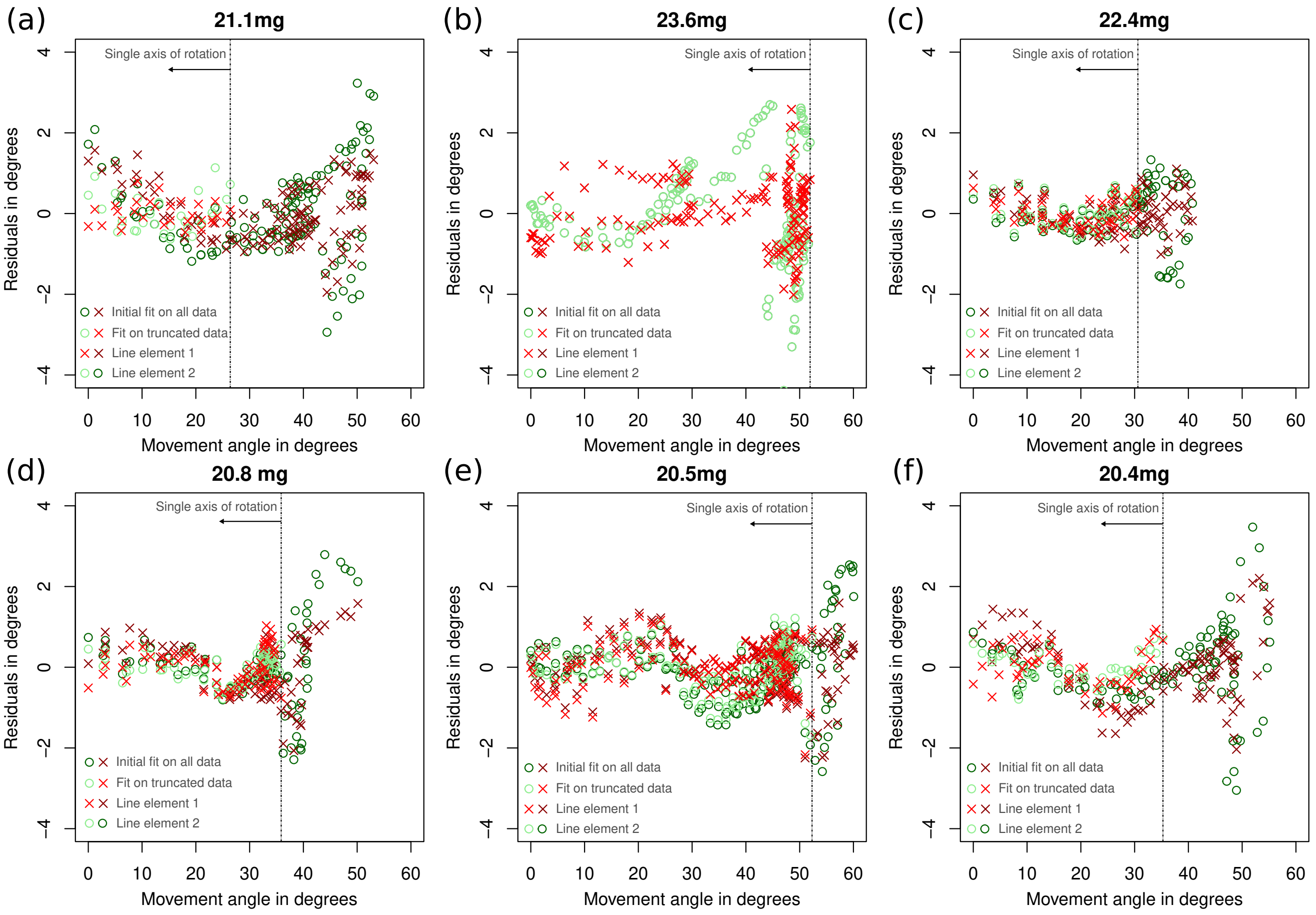


Figure S 2 | **(a-f)** In order to test if the mandibles rotate about a single axis of rotation, we estimated the axis, *bCAR*, that minimises the variation of the angles spanned with two mandible-fixed line elements (see eq. 2.3 in the main manuscript). For each ant, we first calculated *bCAR* across the entire movement angle range, and then calculated the residuals as the difference between expected and observed angle. For large movement angles, corresponding to small opening angles and the region of possible mandible overlap, the residuals were usually higher, suggesting deviation from simple rotation about one fixed axis. This observation was statistically supported by the fact that residuals significantly depended on movement angle and/or linear model assumptions were violated; this held true for all but one ant, see **(b)**. To test if there was a region where the kinematic space could be sufficiently characterised by a single axis of rotation, we successively truncated the data (from large to small movement angles), recalculated *bCAR* and the residuals, and retested both statistical conditions. For each ant, we extracted the largest movement angle range and the minimum opening angle below which the null hypothesis of a single axis of rotation could no longer be statistically rejected (represented by the brighter points. The body mass of each specimen is shown in bold on top of each figure). For further analysis, we used the average minimum opening angle across all ants to divide the mandible kinematic space into two regions: one region in which kinematics can be described via a single axis of rotation, and another in which multiple axes are required.

dépendamment des causes qui peuvent les produire. *Journal de mathématiques pures et appliquées* **5**, 380–440.

**Stokdijk, M., Meskers, C., Veeger, H., De Boer, Y. and Rozing, P.** (1999). Determination of the optimal elbow axis for evaluation of placement of prostheses. *Clinical Biomechanics* **14**, 177–184.

**Woltring, H. J., Huiskes, R., De Lange, A. and Veldpaus, F. E.** (1985). Finite centroid and helical axis estimation from noisy landmark measurements in the study of human joint kinematics. *Journal of Biomechanics* **18**, 379–389.

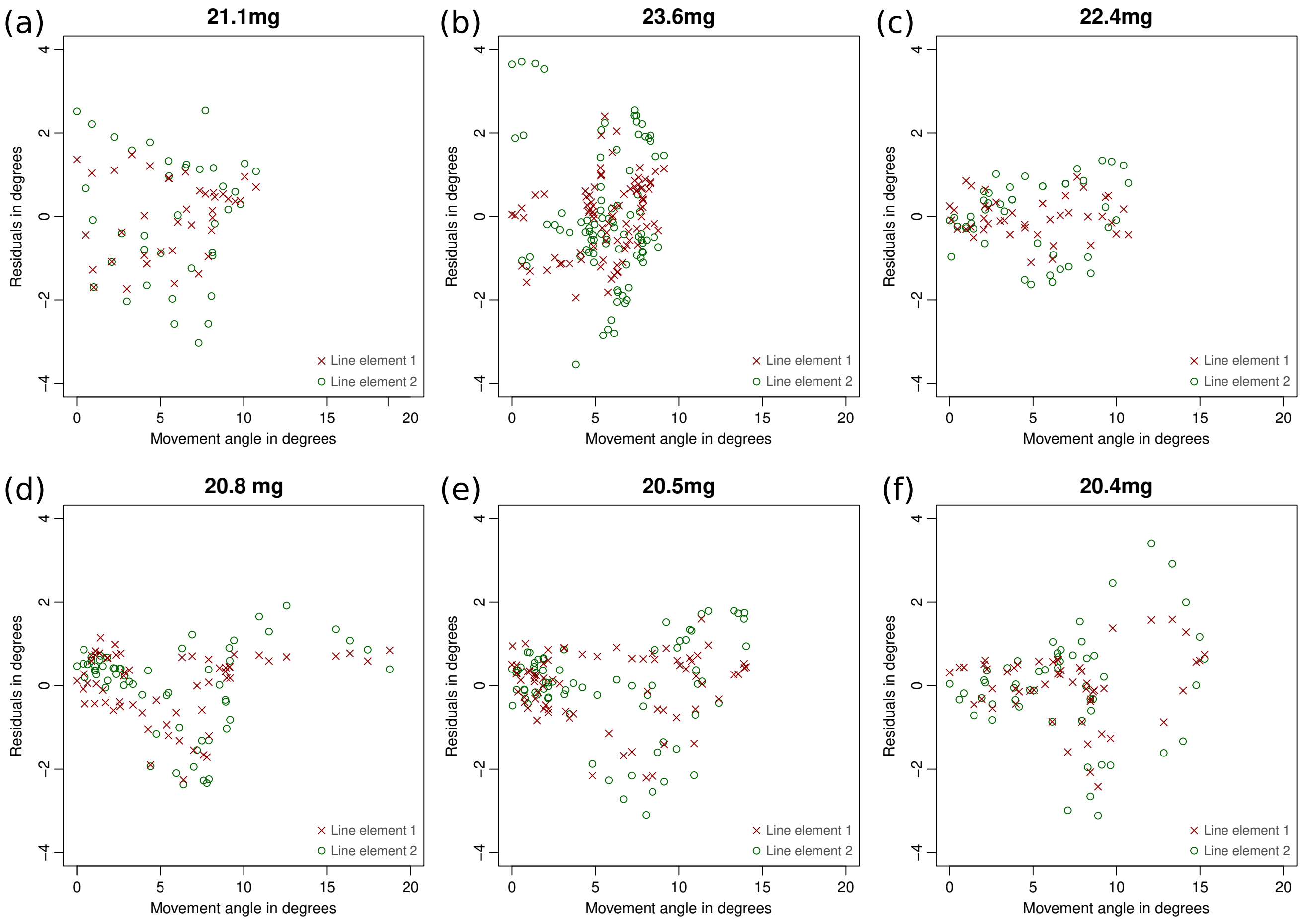


Figure S 3 | **(a-f)** We used statistical arguments to divide the mandible kinematic space into two distinct regions: one region that can be characterised by a single axis of rotation (for opening angles larger than 68°), and a second region where mandibles can interfere with each other and seemingly rotate about at least two axes (for opening angle below 68°). To test statistically if a single axis of rotation indeed fails to adequately describe mandible kinematics in this second region, we identified a dominate axis of rotation on data truncated to opening angles $< 68°$, using eq. 2.3 in the main manuscript. We then calculated residuals and movement angles, performed an ordinary least squares regression on both, and tested if the linear model assumptions were satisfied (for further details, see methods in main manuscript). For all but one ant, **(a)**, the conditions were violated, and the null hypothesis of a single axis of rotation was thus rejected. In further support of this conclusion, the residuals were generally larger than those obtained for the other kinematic region (see Fig. S 2). Note that the movement angles in this figure do not correspond to the angles displayed in Fig. S 2; instead, the smallest movement angles correspond to a maximum opening angle of $\approx 68°$. The body mass of each specimen is shown in bold at the top of each sub-figure.
